# Mind the Bend: Curved and Corrugated Cryo-Lamella for Improved Mechanical Resilience

**DOI:** 10.64898/2026.08.23.746576

**Authors:** Sergey Gorelick, Sylvain Trépout, Patrick Cleeve, Marion Boudes, Yoona Kim, Georg Ramm

## Abstract

Preparing electron-transparent cryo-lamellae is inherently a serial, low-throughput process. During sample handling, milling, and transfer, cryo-fixed cells and their supporting films are subjected to mechanical forces as well as thermal stresses caused by temperature fluctuations. After milling, these extremely thin lamellae remain vulnerable to both mechanical and thermal stress, often leading to cracking or complete disintegration. Consequently, the loss of valuable lamellae is frequently an unavoidable aspect of working with such fragile specimens. In this work, we reconsider the conventional lamella geometry, which is typically a flat, thin cross-sectional slab. During milling, lamellae often become unintentionally bent, complicating the final polishing step required to achieve uniform thinning across their width. To address this limitation, we propose deliberately fabricating lamellae in a pre-bent configuration, i.e. specifically, adopting an arch-shaped profile instead of the traditional flat geometry. The arch shape is intrinsically more mechanically stable than a flat structure, thereby reducing lamella loss due to mechanical failure. Moreover, pre-bent milling patterns facilitate uniform thinning of bent lamellae, which is difficult to achieve using conventional flat milling approaches. In addition to the arch geometry, we investigate corrugated lamellae, characterised by a sinusoidal variation around the plane of a conventional flat lamella. Similarly to the arch shape, the corrugated design offers enhanced mechanical stability compared to traditional flat lamellae. We fabricated a series of test lamellae incorporating both arches and corrugations. High-resolution cryo-TEM imaging was performed to evaluate these structures, demonstrating that non-flat geometries do not compromise cryo-electron tomography performance. Furthermore, finite element method (FEM) simulations were conducted to provide insight into stress distributions within bent and corrugated lamellae.

---

Cryogenic focused ion beam (cryo-FIB) milling is a well-established method for preparing electron-transparent samples for cryo-electron tomography (cryo-ET). In a typical workflow, cells are grown or deposited on electron microscopy (EM) grids, plunge-frozen, and milled directly on the grid to produce thin cryo-lamellae [1, 2, 3, 4, 5, 6]. The resulting lamellae, typically thinner than 200 nm, are transferred to a cryo-transmission electron microscope (cryo-TEM) for high-resolution imaging of cells in a near-native state. Following milling, lamellae are exposed to mechanical and thermal stresses. During transfer between the cryo-FIB and cryo-TEM, temperature fluctuations of several tens of degrees may occur, increasing their susceptibility to damage. Combined with the mechanical forces associated with grid handling, these stresses can destabilise the lamellae, where even minor deformations may lead to cracking or complete failure. Cryo-FIB milling is also inherently low-throughput because lamellae are prepared individually. Each lamella requires approximately 30 minutes of instrument time, excluding sample preparation. Consequently, lamella loss represents a significant loss of time, effort, and instrumentation resources, while also reducing opportunities for cryo-TEM data collection.

Improving the mechanical stability of cryo-FIB-prepared lamellae has therefore become an important research focus. Several approaches have been proposed to increase lamella robustness and survivability. Wang et al. [7] described the fabrication of mechanically stable large-area tissue lamellae and analysed common failure modes (Supplementary Fig. 5d in [7]). Their method disconnects one edge of long lamellae from the bulk material and introduces a “furrow–ridge” design of alternating thin furrows and thicker supporting ridges, effectively creating a “ribbed” or “reinforced” lamella. This approach significantly improves stiffness and reduces bending. Kelly et al. introduced the “notch milling” strategy within the Waffle Method workflow [8, 9]. A ∼200 nm notch partially separates one end of the lamella from the bulk, allowing limited motion and stress relief during handling and transfer. The notch improves lamella survival and facilitates uniform final thinning by reducing stress-induced bending during milling. Wolff et al. developed the “micro-expansion” strategy, in which small trenches are milled on either side of a lamella [10]. Originally developed to reduce grid-induced stress and lamella bending [11], the method was also found to improve survivability.

Reports of lamella failure remain relatively uncommon because published studies generally focus on successful preparations. Nevertheless, available literature [7], community reports [12], and our own observations indicate that fractures most often occur near the attachment points connecting the lamella to the bulk material, suggesting strong local stress concentrations. We previously showed that filleted milling patterns, in which sharp corners are replaced by smooth transitions, reduce stress concentrations and lower the likelihood of lamella failure [13]. More recently, Wachsmuth-Melm et al. [14] proposed a trapezoidal milling geometry. Their results indicate improved mechanical stability due to the smoother transition between the lamella and surrounding bulk material, analogous to corner fillets. Crack formation during thinning, handling, or cryo-TEM imaging can cause catastrophic lamella loss or significant degradation of imaging quality. In our previous study [15], we investigated the incorporation of perforations (crack-arrest holes) into lamellae to improve mechanical stability and mitigate crack propagation.

In the present work, we explore an alternative approach based on *non-planar lamella geometries*. Specifically, we investigate two curved designs: arch-shaped and corrugated lamellae (Fig. 1). Both geometries are expected to be substantially more mechanically robust than conventional flat lamellae. We fabricate these curved lamellae and characterise them using TEM to evaluate their suitability for cryo-ET workflows.

**Figure 1:**
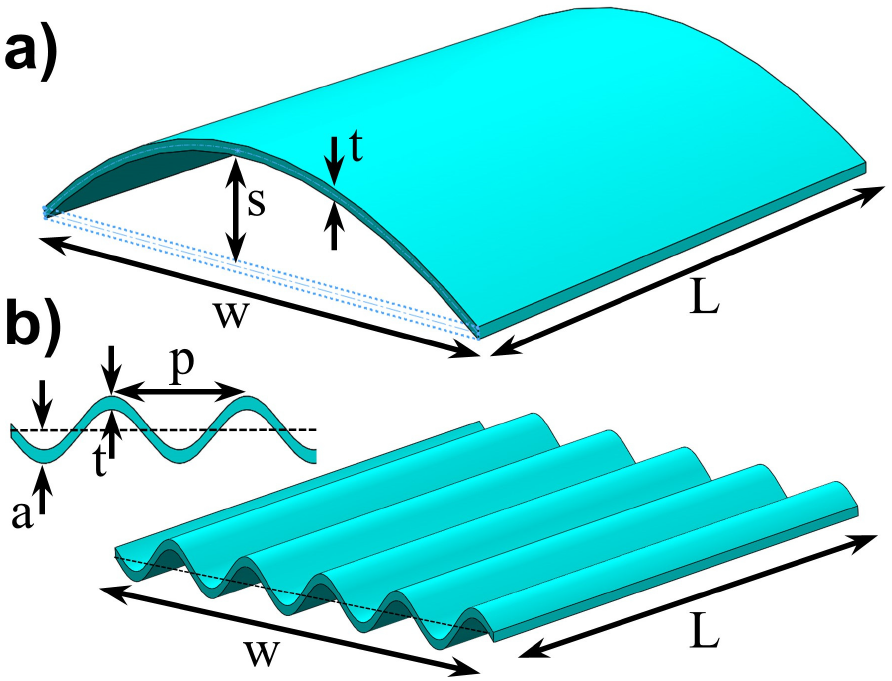
Schematic illustration of the geometric parameters defining (a) an arched lamella with a parabolic profile and (b) a sinusoidally corrugated lamella. (a) Arched lamella. The sag height, *s*, is defined as the distance between the neutral axis of a flat lamella (dashed line) and the neutral axis of the arched lamella. *w* denotes the lamella width, *L* its length, and *t* its thickness. (b) Corrugated lamella. The corrugation pitch, *p*, is the distance between successive corrugation peaks, while the corrugation amplitude, *a*, is defined as the distance between the neutral axis of the corresponding flat lamella and the neutral axis of the corrugated lamella. *t* denotes the lamella thickness.

Cryo-ET workflows are typically based on the assumption that the lamella is flat. In practice, however, this is only an approximation, as lamellae are continuously subjected to mechanical and thermal stresses that can cause them to bend [16] or deform (see Appendix A). A perfectly or nearly flat lamella is inherently mechanically fragile because it offers little resistance to bending and therefore readily deforms in response to external forces. Introducing curvature fundamentally changes the mechanical behaviour of the lamella. Transforming a flat lamella into a parabolically arched structure (Fig. 1a) converts it from a bending-dominated sheet into a shell-like structure. The curvature spreads applied forces across the entire structure, greatly reducing bending and allowing stresses to be carried more uniformly. As a result, an *arched lamella* (Fig. 1a) is substantially more resistant to deformation, buckling, and fracture than a conventional flat lamella of the same thickness. This increased mechanical robustness enables curved lamellae to better withstand the mechanical handling and temperature fluctuations encountered during cryo-FIB preparation, transfer, and cryo-TEM imaging, thereby improving their likelihood of surviving the cryo-ET workflow intact.

Another promising geometry for producing mechanically robust and resilient lamellae is the *corrugated lamella* (Fig. 1b). A corrugated lamella is a geometrically reinforced thin structure that contains the same material and has the same nominal thickness as a conventional flat lamella, yet exhibits substantially different mechanical behaviour. Numerous corrugation geometries exist, including trapezoidal, rectangular, triangular (zigzag), and other profiles. However, for cryo-ET applications, a smooth sinusoidal corrugation is particularly attractive because it can be fabricated more readily and avoids the sharp corners that are associated with local stress concentrations. In a flat lamella, mechanical and thermal loads are accommodated primarily through bending, making the structure highly susceptible to deformation, buckling, and stress concentration. Introducing a sinusoidal corrugation increases the effective stiffness of the lamella and promotes a more uniform distribution of applied loads throughout the structure. As a result, peak stresses are reduced and the lamella becomes more resistant to bending and buckling without requiring an increase in specimen thickness. The corrugated geometry also improves fracture resistance by reducing stress concentrations and hindering crack propagation. Furthermore, thermal expansion and contraction can be partially accommodated through small changes in the corrugation profile. Consequently, corrugated lamellae are expected to be more mechanically stable and more likely to survive cryo-transfers, and cryo-TEM imaging than traditional flat lamellae.

Figure 2 shows an arched lamella milled in yeast cells. Overall, bitmap-based pattern definition proved effective for producing curved lamellae. However, in some specimens the removal of cellular material was less efficient, with residual material occasionally remaining despite repeated ion beam exposure. Based on our experience, the cleaning cross-section milling strategy provides the most reliable material removal, producing cleaner surfaces and leaving fewer residuals than conventional raster scanning approaches. These observations suggest that additional pattern geometries would be a valuable extension of the current lamella-milling toolbox. In particular, curved rectangular milling patterns with a user-defined radius of curvature, combined with cleaning cross-section scan lines that follow the curvature of the lamella, could substantially improve the fabrication of arched structures. Such patterns would also be beneficial for processing lamellae that become bent during milling owing to internal stresses (Appendix A). Uniformly thinning these unintentionally curved lamellae using conventional straight milling patterns is often challenging, and typically only a small portion of a thick bent lamella can be polished to the desired thickness. Curved milling patterns that match the lamella geometry would therefore facilitate more uniform thinning and improve the yield of usable lamellae.

**Figure 2:**
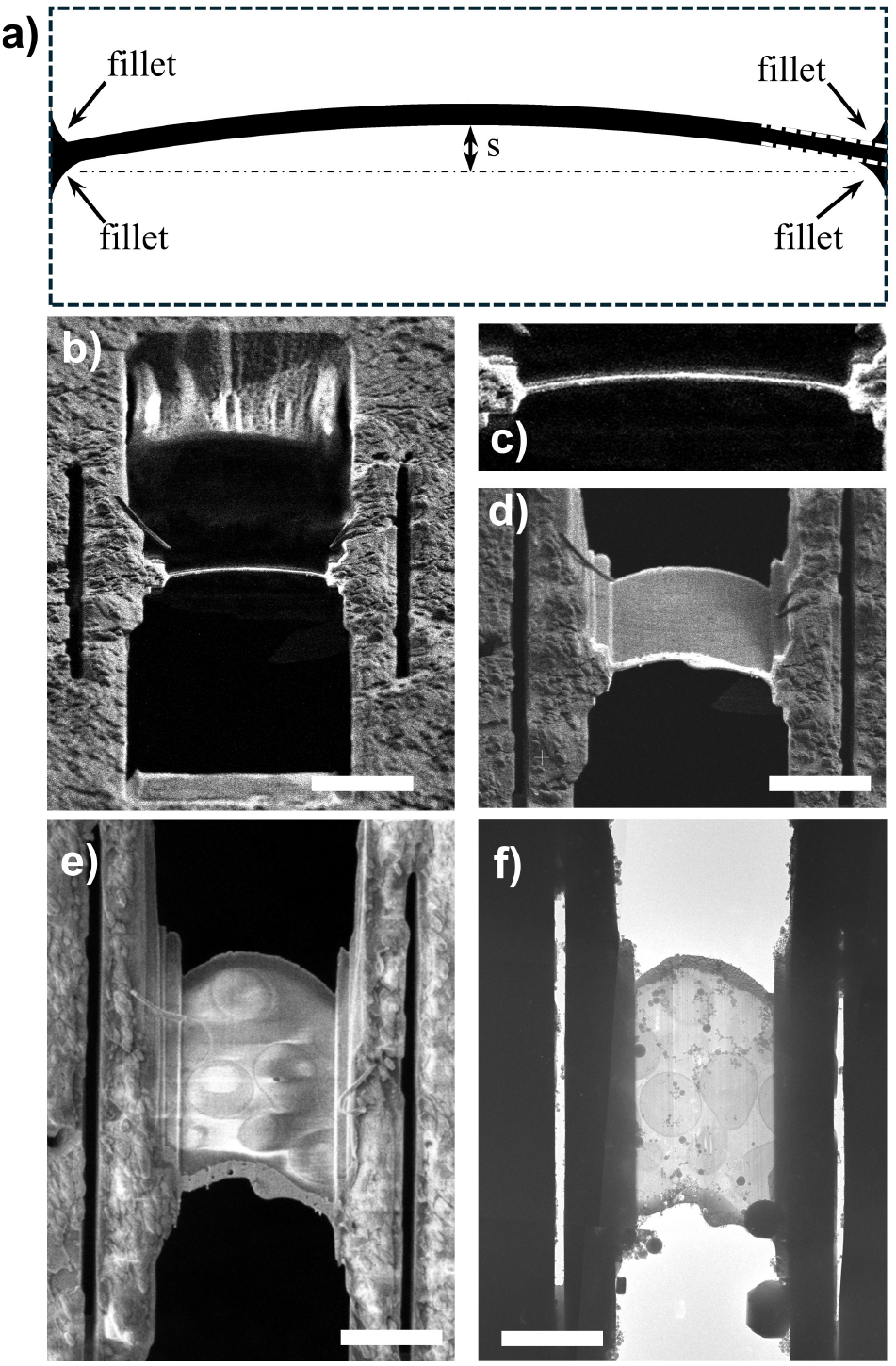
Bent, arched lamella by design. (a) Bitmap milling pattern generated using a custom script. The pattern is defined by parameters including the sag height *s*, width, and thickness of the parabolically arched lamella. Corner fillets are incorporated to smoothen sharp transitions and reduce stress concentrations. The dashed outline indicates the extent of the bitmap pattern. White pixels correspond to regions exposed to the ion beam and milled away, whereas black pixels indicate regions that remain unexposed and therefore unmilled. (b) FIB image of an arched lamella prepared in a yeast cells, viewed along the milling direction. The lamella has a sag height of 400 nm. (c) Magnified view of the arched lamella in (b). (d) FIB image of the same arched lamella after tilting the sample by 18°. (e) SEM image of the lamella viewed from the milling-angle direction. (f) Low-magnification TEM image (3600×) of the arched lamella. Scale bar is 5 *µ*m.

In this study, we fabricated arched lamellae with sag heights ranging from 50 nm to 500 nm, corresponding to up to approximately 2.5 times the thickness of a typical lamella. In general, lamellae with an upward arch were easier to fabricate, whereas downward-arched lamellae proved more challenging, often resulting in over-milling and reduced control over the final geometry (Appendix B). From our experience, lamellae tend to bend upward more frequently than downward (Appendix A). Whether this behaviour reflects an inherent mechanical property of the lamella–grid system or is a consequence of the preparation workflow [17, 18] remains unclear. In our milling procedure, the upper side of the cell is milled first, followed by the lower side adjacent to the grid, and this sequence may contribute to the observed bending. The origin of this behaviour remains an open question and will be investigated in future work.

A flat lamella behaves mechanically like a thin beam or plate and therefore responds to external loads primarily through bending. In contrast, a curved structure redistributes applied loads across its entire surface through a combination of tensile and compressive stresses, greatly reducing the extent of bending. As a result, even a modest degree of curvature (or a small sag) can significantly improve the mechanical robustness of a lamella. An arched lamella therefore begins to outperform a flat lamella in resistance to deformation, buckling, and fracture even at relatively small radii of curvature. The greater load-bearing capacity of an arched lamella offers several potentially important advantages for cryo-EM applications.

Firstly, because a curved lamella is inherently stronger than an equivalent flat lamella, it may be possible to fabricate thinner lamellae without compromising mechanical stability. Thinner lamellae are desirable in cryo-EM because they reduce electron scattering and can improve image quality and achievable resolution.

Secondly, the increased structural robustness of an arched lamella may allow the fabrication of wider lamellae while maintaining mechanical stability comparable to that of narrower flat lamellae. This would increase the usable imaging area and potentially enable the collection of more structural information from a single lamella.

Another advantage of an arched lamella is that its curved geometry increases the effective imaging volume. This increased volume of electron-transparent material enables the acquisition of more tomograms and the observation of a greater number of cellular features within a single lamella, thereby increasing the amount of structural information that can be collected from each sample.

The mechanical behaviour of arches and curved shells is well established in structural mechanics [19, 20, 21]. In contrast to flat plates, which primarily resist external loads through bending, curved structures can redistribute loads through a combination of bending, tension, and compression, resulting in greater stiffness and improved resistance to deformation. Classical shell theory predicts that the stiffness of an arch increases with increasing curvature (or sag), allowing substantially larger loads to be carried without a corresponding increase in material thickness. For shallow arches, the ratio between the stiffness of an arched structure and that of a corresponding flat beam or plate can be expressed as a function of the arch geometry. For example, beam and shell theory can be made to show (for a parabolic beam, with fixed-fixed boundary condition, see also Appendix C for simulated results) an increase in effective stiffness according to *k*_arch_ ≈ *k*_flat_ (1 + (*s/t*)^2^), where *k*_flat_ and *k*_arch_ are the stiffnesses of the flat and arched structures, respectively, *s* is the arch sag, *t* is the lamella thickness. Although simplified analytical models provide useful insight, the behaviour of cryo-lamellae is more accurately captured using finite element method (FEM) simulations, which account for the full three-dimensional geometry and loading conditions. Figure C.7a shows finite element methood (FEM) simulations of an arched lamella subjected to a uniform load. In the present case, the effective spring constant obtained from FEM under a uniformly distributed load cannot be directly interpreted as a complete measure of mechanical robustness, since the stress distributions and deformation modes differ substantially between flat and arched geometries. Nevertheless, it provides a useful comparative metric. FEM analysis of a 10 *µ*m × 10 *µ*m × 200 nm lamella showed that a flat lamella is approximately ten times more compliant than a corresponding arched lamella with a sag of 600 nm, and approximately five times more compliant than an arched lamella with a sag of 400 nm. These results demonstrate that even modest curvature produces a substantial increase in structural stiffness and resistance to deformation.

Figure 3 shows two sinusoidally corrugated lamella milled in yeast cells. Overall, bitmap-based pattern definition proved effective for producing curved lamellae. Corrugated structures are among the most widely used forms of geometric reinforcement and are found in applications ranging from corrugated cardboard packaging to roofing panels and large structural components [22, 23, 24, 25]. Their key advantage is that they provide substantially greater robustness and mechanical stiffness to bending across the corrugations without requiring additional material thickness. Applied to cryo-lamellae, this means that a corrugated lamella can potentially be made thinner than a conventional flat lamella while maintaining comparable mechanical stability, or alternatively, made wider without compromising its resistance to deformation and fracture. We also evaluated stepped (terraced) lamellae, which were found to be straight-forward to fabricate and mechanically viable (Appendix D). Unlike arched or corrugated geometries, terraced lamellae can be produced without bitmap patterns by using a series of progressively offset cleaning cross-section polishing windows. In the present study, we focused primarily on sinusoidally corrugated lamellae. A range of corrugation pitches, from 500 nm to 2 *µ*m, was investigated while maintaining a constant pitch-to-amplitude ratio of 10 (e.g,, a corrugation pitch of 2 *µ*m corresponded to an amplitude of 200 nm). Figure C.7b shows finite element simulations of a corrugated lamella subjected to a bending moment. The corrugated lamella is an anisotropic structure whose mechanical response depends on the loading direction. The corrugations increase the effective bending stiffness and improve resistance to local buckling and deformation, particularly for loads applied perpendicular to the corrugation direction. Unlike an arched lamella, which primarily enhances global structural rigidity by redistributing loads across the entire lamella, corrugations provide local geometric reinforcement and redistribute stresses over a series of peaks and valleys. This reduces peak stress concentrations and results in a more uniform stress distribution throughout the structure. Consequently, corrugated lamellae may be particularly advantageous in situations involving localised mechanical loads, thermal gradients, or stress concentrations. In addition, the corrugations can accommodate part of the applied deformation through local changes in shape, allowing the structure to absorb and redistribute stress more effectively than a flat lamella. As a result, corrugated lamellae are expected to exhibit improved resistance to crack initiation and propagation, thereby enhancing their overall mechanical robustness and survivability. Similarly to arched lamellae, corrugation also increases the overall lamella length, but the effect is substantially more pronounced (Appendix C). As the corrugation amplitude increases, the arc length of the lamella grows significantly beyond its projected width, resulting in a larger electron-transparent volume within the same footprint. Consequently, corrugated lamellae provide a more effective means of increasing the amount of imaged material and the volume available for cryo-EM data collection than arched lamellae.

**Figure 3:**
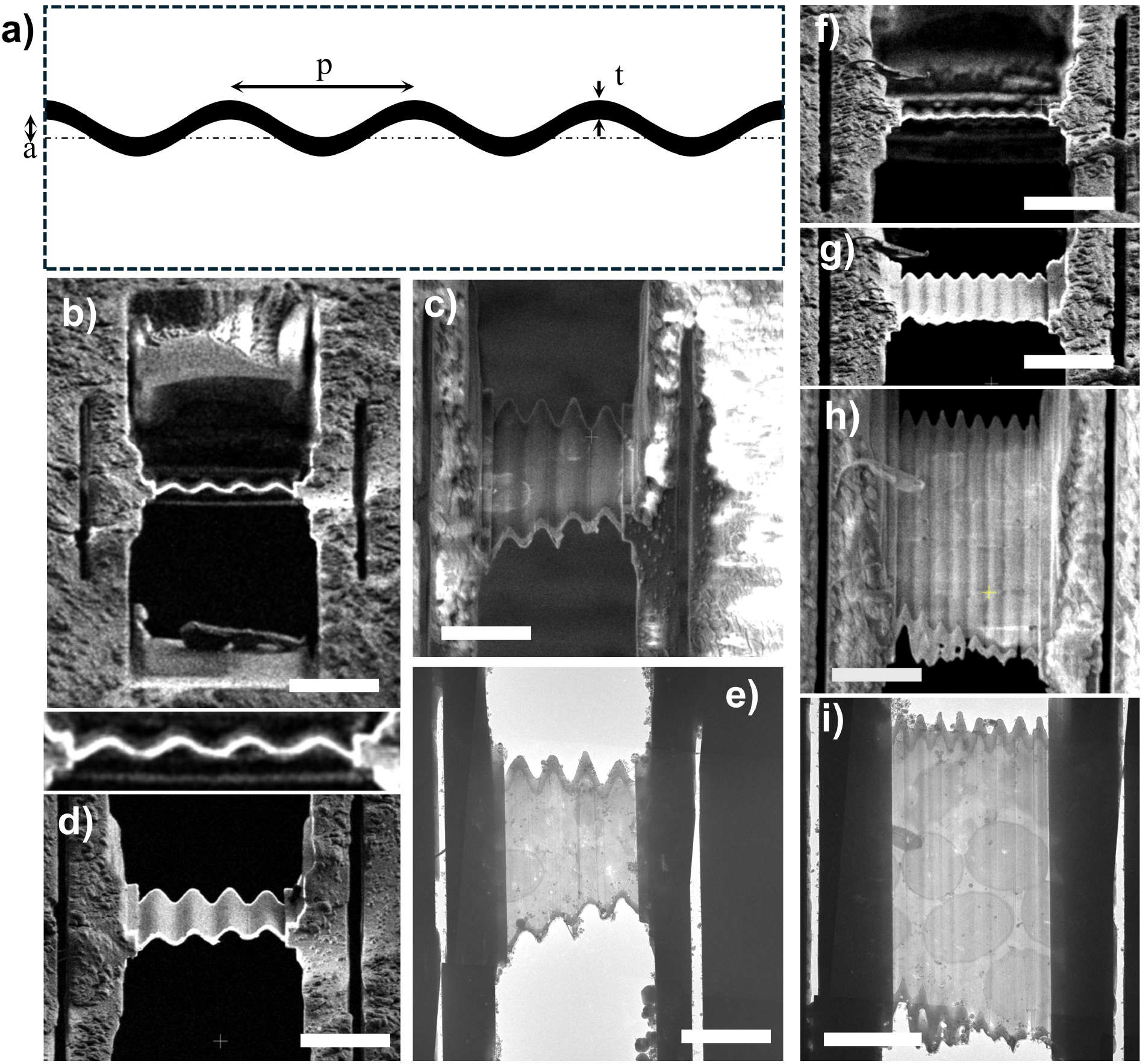
Corrugated sinusoidal lamellae. (a) Bitmap milling pattern generated using a custom script. The pattern is defined by the corrugation pitch *p*, corrugation amplitude *a*, and lamella thickness *t*. The dashed outline indicates the extent of the bitmap pattern. White pixels correspond to regions exposed to the ion beam and milled away, whereas black pixels indicate regions that remain unexposed and therefore unmilled. (b) FIB image of a corrugated lamella prepared in a yeast cell, viewed along the milling direction. The corrugation pitch is 2 *µ*m and the amplitude is 200 nm. (c) SEM image of the lamella shown in (b), viewed from the milling-angle direction. (d) FIB image of the lamella shown in (b) after tilting the sample by 13°. (e) Low-magnification TEM image (3600×) of the corrugated lamella with a 2 *µ*m pitch and 200 nm amplitude. (f) FIB image of a second corrugated lamella prepared in a yeast cell, viewed along the milling direction. The corrugation pitch is 1 *µ*m and the amplitude is 100 nm. (g) FIB image of the lamella in (f) after tilting the sample by 8°. (h) SEM image of the lamella shown in (f), viewed from the milling-angle direction. (i) Low-magnification TEM image (3600×) of the corrugated lamella in (f,h). Scale bar is 5 *µ*m.

To date, we have fabricated 18 arched and 16 corrugated lamellae, none of which failed during cryo-transfer. While the sample size is currently too small to support statistically significant conclusions, we plan to expand this dataset in future studies to enable a more rigorous assessment of lamella survivability and transfer reliability.

One might expect that arched and corrugated lamellae are unsuitable for cryo-ET because conventional workflows assume a flat specimen. In reality, however, lamellae are often already bent or deformed due to preparation and residual stresses [16], so perfect flatness is rarely achieved. We successfully collected tomograms (see Fig. 4) from all tested arched and corrugated lamellae, including those with the largest sag and corrugation amplitude values, without any imaging difficulties. The maximum sag in this study was approximately 500 nm (maximum of only 200 nm corrugation amplitude or 400 nm peak-to-valey), whereas typical TEM defocus values were 1.5–2.5 *µ*m. Thus, the height variation introduced by the arch or corrugation was only a small fraction of the applied defocus and did not noticeably affect data acquisition or reconstruction [26]. These results demonstrate that moderately arched lamellae remain fully compatible with cryo-ET while providing substantially improved mechanical robustness compared with conventional flat lamellae.

**Figure 4:**
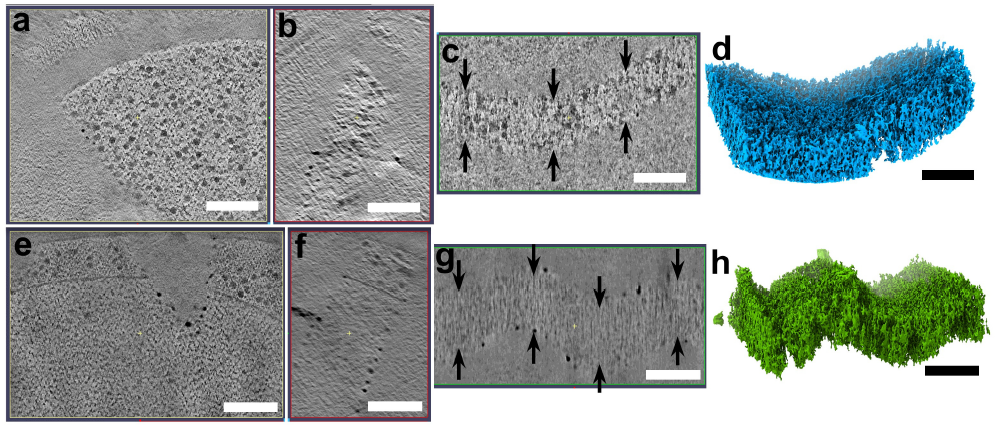
Cryo-electron tomography and volume reconstruction of (a–d) a curved lamella with a sag of 500 nm and (e–h) a corrugated lamella with a pitch of 500 nm and corrugation amplitude of 50 nm. Panels (a,e), (b,f), and (c,g) show orthogonal projections of the reconstructed tomographic volumes, while (d,h) show the corresponding three-dimensional volume renderings. Scale bar is 200 nm.

In summary, arched and corrugated lamellae improve mechanical robustness through different mechanisms. An arched lamella introduces curvature across the entire structure, allowing loads to be distributed through tensile and compressive stresses rather than primarily through bending. This shell-like behaviour increases resistance to large-scale deformation, buckling, and fracture. In contrast, a corrugated lamella uses a periodic wave-like geometry that increases effective stiffness and distributes stresses more uniformly without increasing lamella thickness. The corrugations act as built-in reinforcement features that reduce local bending, improve resistance to crack initiation, and help accommodate thermal stresses. Consequently, the arch primarily enhances global mechanical stability, whereas the corrugations improve local mechanical stability. A lamella combining both geometries could potentially benefit from the advantages of each, providing both improved overall rigidity and enhanced resistance to localised deformation and failure.

## Materials and methods

### Cell culture on EM grids and freezing

Saccharomyces cerevisiae strain derived from BY4741 was grown overnight in 5 mL YPD medium (1% yeast extract, 2% peptone, 2% glucose) at 30 °C, 160 rpm. The culture was then kept at 4 °C until use. 5 *µ*L of the cell-containing solution was deposited on a Cu-200 R2/2 grid (Quantifoil™) which had previously been glow-discharged for 30 s using a Pelco EasyGlow™. Excess solution was removed through manual blotting. The cells were vitrified by plunge freezing in a liquid 60/40 Ethane/propane mix and stored in liquid nitrogen.

### Cryo-FIB milling

Lamellae fabrication was performed on a dual-beam ThermoFisher Scientific (TFS) Helios 5 UX Cryo-FIBSEM using 30 keV Ga+ beam. The grids clipped in autogrids were cryogenically transferred into the cryo-FIB, and suitable cells for thinning and milling were selected from mapping and imaging the grids with electron and ion beams. The microscope is equipped with a cryogenic shuttle with a pre-tilt of 40°. The ion beam incidence with respect to the grid plane was typically set either to 12 or to 15° unless specified otherwise (stage tilt 14° or 17°), and kept constant throughout the lamella milling. We ensured that each selected milling site was at the eucentric point of the microscope, *e*.*g*., the point of coincidence between the electron and ion beams. In this point the sample is at 4 mm from the electron microscope’s polepiece. Next, the sample was lowered to 8 mm from the electron polepiece and a layer of organometallic platinum was deposited onto the grids using gas injection system (GIS) operated at 29 °C for 6 s. The lamellae milling was aided with AutoLamella (fibsemOS) software [27, 28]. Firstly, we translated the stage from one lamella site to the next, milling fiducial markers at each site to facilitate future alignments. The fiducial marks were milled using Rectangle milling pattern. Next, we proceeded to each lamella site, aligning using the previously milled fiducial markers.

Before the rough milling, we created micro-expansion gaps using a 2.6 nA beam current. The microexpansion gaps were milled using Rectangle milling pattern. The microexpansion gaps were positioned on either side of the lamella 10 *µ*m from its centre.

We then performed rough and intermediate milling with beam currents of 2.6 nA and 0.44 nA, respectively, to define the initial, relatively thick lamellae. The rough milling step aimed at creating a 5-*µ*m-thick and 12-*µ*m-wide lamella, while the intermediate milling step aimed at creating a 1.5-*µ*m-thick and 11-*µ*m-wide lamella. Rough and intermediate milling steps were performed using “cleaning cross-section” patterns.

The final thinning of the thick lamellae was carried out sequentially using a 90 pA ion beam, producing lamellae with a nominal thickness of 180 nm. To fabricate curved and corrugated lamellae, we generated custom bitmap milling patterns that selectively removed material while preserving the desired arch or sinusoidal corrugated profile. In these bitmap masks, pixels with a value of 255 (white) corresponded to regions exposed to the ion beam and milled away, whereas pixels with a value of 0 (black) were skipped by the beam and therefore remained intact. This approach enables the fabrication of complex binary patterns and even three-dimensional surface profiles [29].

We initially tested a bitmap pixel size of 5 nm, however, discretisation artefacts were clearly visible in the resulting lamella surfaces. Reducing the pixel size to 2 nm produced substantially smoother and more uniformly polished lamellae. A pixel dwell time of 3 *µ*s was used, as shorter dwell times resulted in inefficient material removal and left residual material that persisted even after prolonged beam exposure. Increasing the dwell time improved both material clearing and the overall milling rate, yielding cleaner and more reproducible lamella surfaces.

The Autolamella has recorded the process and intermediate steps in images. All SEM images were collected at 2 keV beam energy, 100 pA beam current, 1 *µ*s dwell time and 1536×1024 pixels image resolution. All FIB images were acquired at 30 keV beam energy, 41 pA beam current, 1 *µ*s dwell time and 1536×1024 pixels image resolution.

### Cryo-TEM tomography

Freshly milled lamellae were loaded into a Titan Krios G4 (ThermoFischer Scientific) cryo-TEM to avoid ice contamination (loading within 12 hours after fine milling). The Titan Krios G4 was operated at 300 kV, and low-dose images were collected on a Falcon 4i direct electron camera post Selectris-X energy filter. The images of the cryo-FIB lamellae presented in this work were recorded using the TFS Tomography TOMO5 software at two different magnifications, 740 and 3,600×. Tilt-series were also recorded using the TOMO5 software. Tilt-images were collected at 53,000× magnification (corresponding pixel size at the sample level was 2.39 Å), using a 20 eV slit, and saved as Electron-Event Representation (EER). A dose-symmetric data collection scheme was used [30], collecting images up to ±56° around the angle at which lamellae were milled (either +12 or +15°), using 3° tilt increments. The total electron fluence used per tilt-series was 100-120 e^−^/Å^2^.

### Cryo-tomography analysis

Motion-correction and CTF estimation were performed in Warp [31], outputting even/odd image pairs for further denoising. Alignment of the tilt-series was performed using patch tracking from the Imod wrapper of Warp [32, 33]. Further refinement of the tilt-series alignment was performed using MissAlignment [34], and reconstruction of the even/odd volume pairs was computed in Warp. Denoising and missing-wedge correction were performed using IsoNet2 [35]. Reconstructed tomograms were investigated and figures were generated using Imod [32, 33] and ChimeraX [36].

## Acknowledgement

The authors acknowledge the use of instruments and assistance at the Monash Ramaciotti Centre for Cryo-Electron Microscopy, a Node of Microscopy Australia. This research used equipment funded by Australian Research Council grants: FEI Helios Cryo FIBSEM - ARC LIEF (LE150100132) and Titan Krios - ARC LIEF (LE120100090). This research was in part funded by ARC DP200103637. This work has been made possible in part by CZI grant DAF2021-225399, grant DOI https://doi.org/10.37921/334038myxhsa (to GR) and grant 2025-366327 (5022) GB-1633987 (to PC & GR) from the Chan Zuckerberg Initiative DAF, an advised fund of Silicon Valley Community Foundation (funder DOI: 10.13039/100014989).

## Appendix A. Lamellae bent during or after FIB milling

**Figure A.5:**
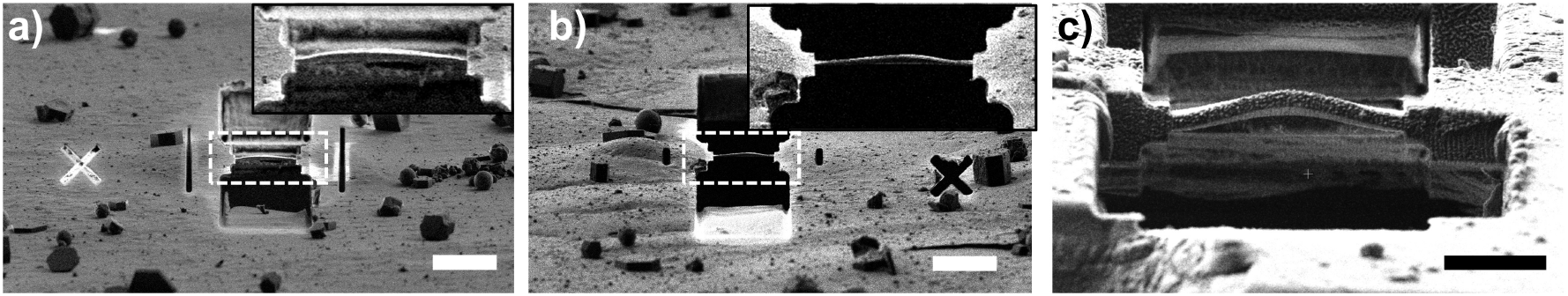
Cryo-lamellae bent during or after FIB milling. (a,b) FIB images acquired from the lamella-milling direction. The 9.5 *µ*m wide lamellae were prepared within cells grown on EM grids [13] Following the final polishing step, the lamellae exhibited pronounced upward bending (with the downward direction corresponding to the side of the cell adjacent to the grid). Insets show magnified views of the lamellae, highlighting the upward curvature. (c) Large Waffle-method lamella approximately 23 *µ*m in width and 1.5 *µ*m in thickness after the rough-milling stage. The lamella displays substantial upward bending, likely caused by the release of residual stresses within the surrounding Waffle material during milling. Scale bar is 10 *µ*m.

## Appendix B. Arched lamella, downward bending

**Figure B.6:**
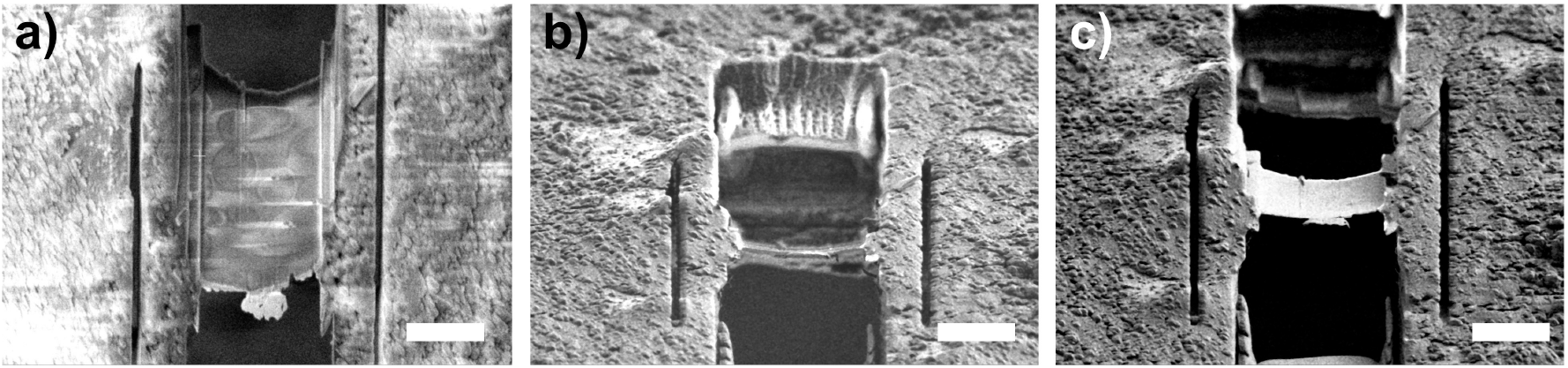
Arched cryo-lamella made by FIB milling with the downward bending. (a) SEM image of 8 *µ*m wide lamellae were prepared within yeast cells on an EM grid. (b) FIB image of lamella in (a) after the final thinning viewed from the milling-angle orientation. (c) FIB image of the same lamella shown in (b) after tilting the sample by 8°. Scale bar is 5 *µ*m.

## Appendix C. FEM Simulations

For the mechanical analysis, we model the lamella as a plate with the width *w* of 10 *µ*m, length *L* of 10 *µ*m, and thickness *t* of 0.2 *µ*m and compare it with a corresponding arched lamella of identical width, length, and thickness, with the sag height *s* treated as a variable parameter. For a shallow parabolic arch, the radius of curvature *R*_*c*_ can be expressed as a function of the arch sag and span/width according to 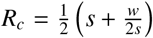 . Given the lamella’s small thickness relative to its lateral dimensions, the flat lamella can, to a good approximation, be treated as a thin beam or plate. We assume fixed–fixed boundary conditions, meaning that the two opposite edges of the flat and arched structures are constrained against displacement. For the special case of a uniformly distributed load acting on the structure, analytical solutions are available for the deflection of a fixed–fixed beam. For a beam subjected to bending, the second moment of area is given by the following expression:

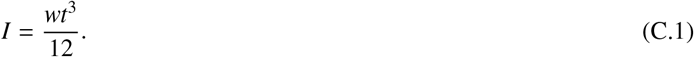

Because the lamella thickness is small, and the second moment of area scales with the cube of the thickness *t*^3^, its bending stiffness is correspondingly very low. As a result, the lamella is highly compliant and readily deforms under relatively small mechanical loads. For the lamella geometry considered here, the second moment of area is only 6.67×10^−27^ m^4^. For a uniformly distributed load *F* acting along the entire beam length, the maximum deflection at the beam centre can be expressed as

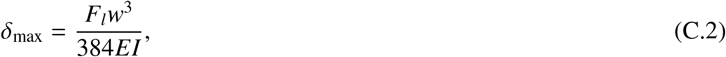

where *E*=1 GPa is the Young’s modulus of vitrified ice[13], *F*_*l*_ = *F/w* is the force on the beam per unit of its width. For a uniform load of 1 *µ*N, and hence the force per length of 0.1 N m^−1^ for *w*=10 *µ*m, the maximum displacement at the lamella centre is 392 nm. This result shows excellent agreement with the displacement predicted by FEM simulations (Fig. C.7a). The spring constant, *k*_flat_ = *F/δ*_max_ is therefore 2.56 N m^−1^. Table C.1 compares the effective spring constants of arched lamellae as a function of arch sag. The corresponding increase in lamella length arising from the introduction of curvature is reported as well. As shown in Table C.1, the stiffness ratios are in excellent agreement with the simplified theoretical relation *k*_arch_ ≈ *k*_flat_ (1 + (*s/t*)^2^).

**Table C.1:**
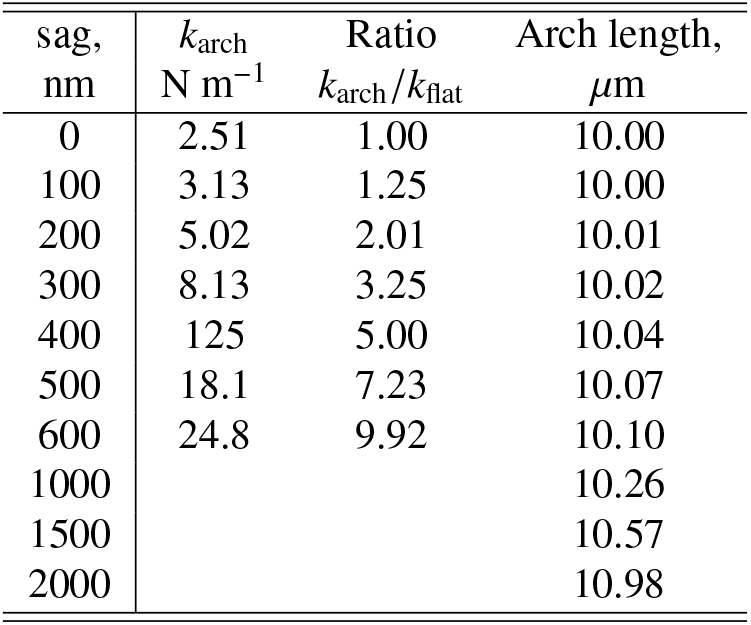
Spring constant of an arched lamella with *w*=10 *µ*m, *L*=10 *µ*m and *t*=0.2 *µ*m for different sags *s*. The values are derived from FEM simulations. The lamella is rigidly fixed at opposite edges and a uniform load is assumed. Ratio of the spring constant shown with respect to the corresponding flat lamella, and increase in arch length is shown for select sags.

**Figure C.7:**
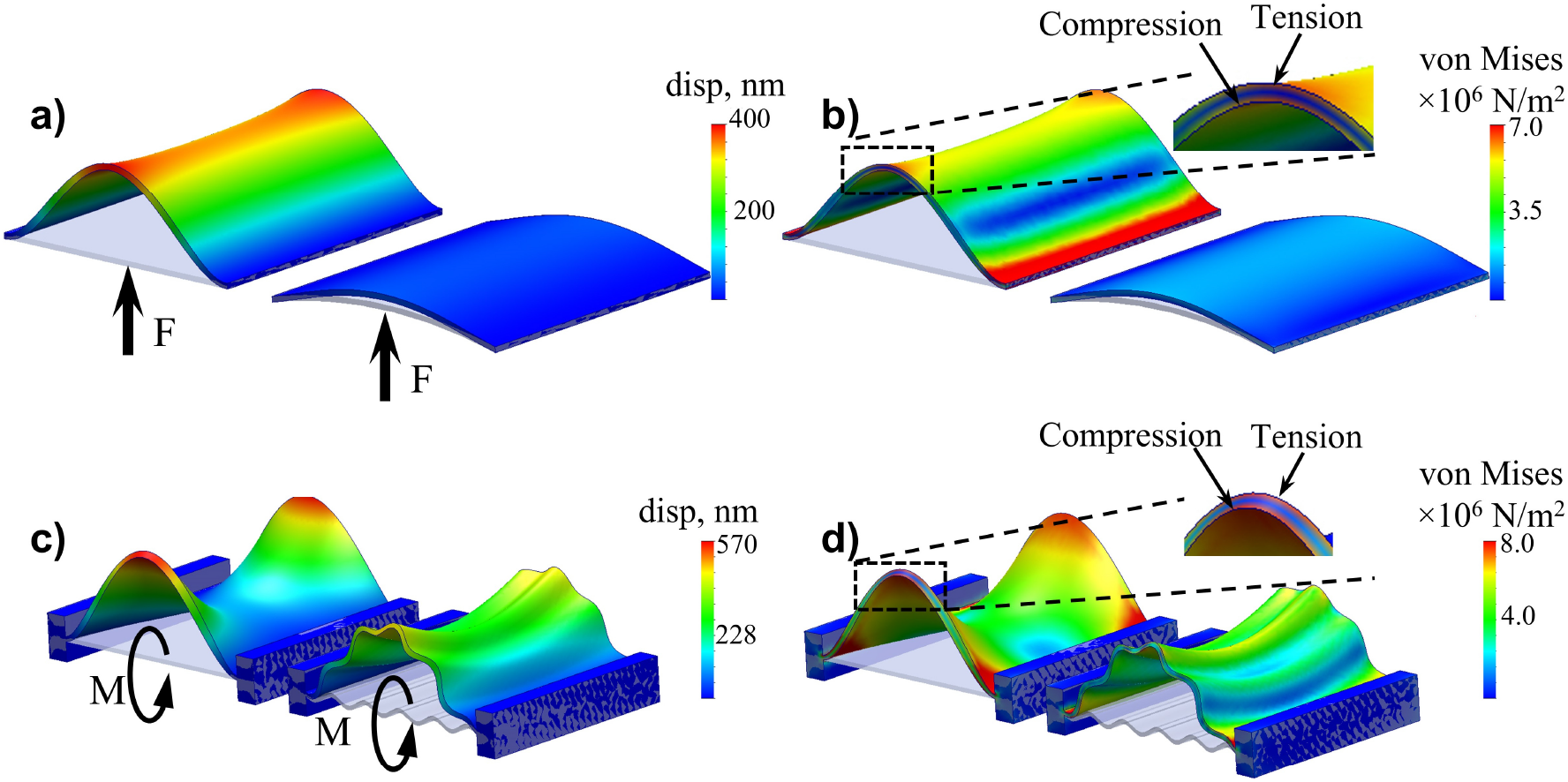
(a,b) Comparison of deformation and von Mises stress, respectively, in a flat lamella and an arched lamella with a sag of 600 nm subjected to a uniformly distributed load, *F*, of 1 *µ*N. The magnified inset in (b) illustrates the characteristic deformation of a bending beam, with one side in compression and the opposite side in tension. In contrast, the arched lamella exhibits substantially smaller deformation and a more uniform stress distribution throughout the structure. (c,d) Comparison of deformation and von Mises stress, respectively, in a flat lamella and a corrugated lamella with a corrugation pitch of 2 *µ*m and an amplitude of 200 nm subjected to a total bending moment *M* of 10 *µ*N m^−1^ applied at the opposite edges. The magnified inset in (d) highlights the typical stress distribution in the flat lamella, where bending produces compression on one side and tension on the other. By comparison, the corrugated lamella undergoes considerably less deformation and displays a more homogeneous distribution of stress across the entire structure. The deformations are exaggerated by a factor of 8×.

Figure C.7b shows finite element simulations of a corrugated lamella subjected to a bending moment. In a flat lamella, bending produces compression on one side and tension on the other, leading to localised stress concentrations, whereas the corrugated lamella undergoes markedly less deformation and exhibits a more homogeneous stress distribution across the entire structure. Unlike an arched lamella, which primarily improves global structural rigidity, corrugations provide local geometric reinforcement, allowing stresses to be redistributed and partially accommodated through local shape changes, thereby improving resistance to deformation, crack initiation, and crack propagation.

For corrugated lamellae, the increase in lamella length is substantially greater than for arched lamellae. Determining the exact length of a sinusoidal profile is not straightforward analytically, particularly for larger amplitudes. However, it can be readily evaluated using CAD software by directly measuring the length of the generated curve For a corrugation pitch of 2 *µ*m and an amplitude of 100 nm, the lamella length increases from 10 *µ*m to 10.24 *µ*m. As the corrugation amplitude increases, the lamella length grows rapidly: a 200 nm amplitude produces a length of 10.92 *µ*m, a 400 nm amplitude increases the length to 13.19 *µ*m, and a 600 nm amplitude results in a length of 16.17 *µ*m. Thus, corrugation significantly increases the surface area and effective imaging volume of the lamella while maintaining the same projected width and nominal thickness.

## Appendix D. Lamella with terraced/stepped corrugation

**Figure D.8:**
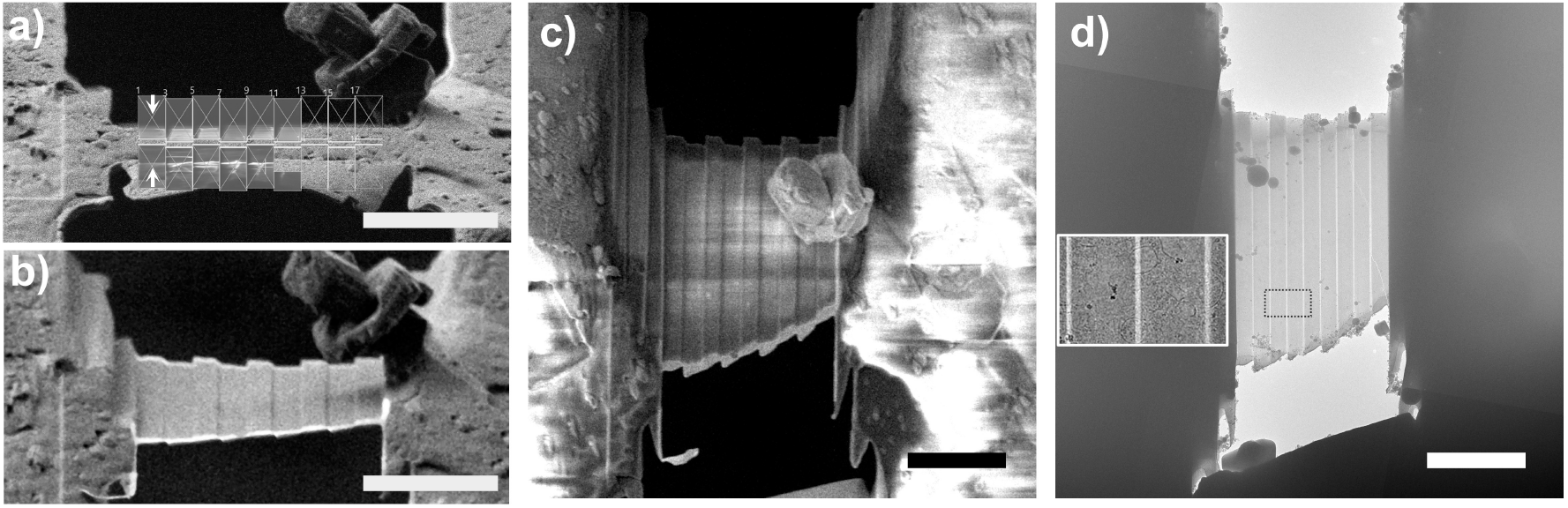
Terraced corrugated lamella. (a) FIB image of a cell during milling of a terraced corrugated lamella. The terrace pitch is 2 *µ*m and the corrugation amplitude is 100 nm. The polishing region of a conventional flat lamella was subdivided into a series of 1 *µ*m-wide cleaning cross-section patterns, with each successive segment offset by ±100 nm in the vertical direction to generate the stepped profile. (b) FIB image of the lamella in (a) after tilting the sample by 10°.(c) SEM image of the fabricated lamella showing the terraced corrugation geometry. (d) Low-magnification TEM image (3600×) of the terraced lamella with a 2 *µ*m pitch and 100 nm amplitude. The inset shows a higher-magnification view of cellular features within the lamella, with the thin transition regions between adjacent terraces at slightly different heights clearly visible. Scale bar is 5 *µ*m.

